# Synthetic Hydrogels with Entangled Neutrophil Extracellular Traps Influence Tumor Progression in MDA-MB-231 Cells

**DOI:** 10.1101/2023.09.27.559781

**Authors:** Rachel R. Katz, Shamitha Shetty, Jennifer L. West

## Abstract

We incorporated neutrophil extracellular traps (NETs) in a poly(ethylene glycol)-based synthetic extracellular matrix to study their impact on tumorigenesis in triple negative breast carcinoma (TNBC) cells in a highly controlled environment. We observed that NETs helped to increase cell survival, proliferation, and pro-metastatic morphological phenotype. We also showed that the presence of NETs influenced the secretion of IL-8, a pro-NETosis factor, and that conditioned media from cells cultured in these gels influenced NETosis in an IL-8 dependent manner. The results observed in this system correlate with murine models and clinical studies in the literature and help to provide additional insight of the individual factors at play in the NET-mediated promotion of TNBC progression and metastasis.

## Introduction

Neutrophils, a subset of polymorphonuclear immune cells, are the most abundant leukocyte in human blood [1, 2]. They are the first infiltrating cells in the innate immune response, participating in the entrapment and clearance of pathogens. One of their key roles is NETosis, or the process by which neutrophils decondense their chromatin structure, expelling a web of DNA complexed with granular enzymes like myeloperoxidase (MPO) called a neutrophil extracellular trap (NET), thus entrapping a target. In the case of non-lytic NETosis, this process results in a phagocytic cytoplast which can engulf pathogens. In lytic NETosis, the neutrophil membrane is ruptured, and other immune cells such as macrophages are required for debris clearance [1, 2]. Classical NETosis is triggered in response to numerous stimuli, including reactive oxygen species (ROS), bacterial lipopolysaccharides (LPS), immune complexes, and inflammatory cytokines [2, 3].

As better mechanisms for detection have arisen, our understanding of NETs in disease has dramatically increased in the last decade. In cancer biology, neutrophils and NETosis, like many other aspects of the immune system, may play a dual anti-tumoral/pro-tumoral role depending on the stage of disease [2, 4–6]. Early in disease development, neutrophils are more likely to be found around rather than within the tumor [6]. Thus, NETosis in tumor-surveilling tumor-associated neutrophils (TANs) may act like NETosis in the presence of pathogens: a way to trap tumor cells, preventing migration and aiding surveillance by other immune cells, and exerting cytotoxicity via granular enzymes [7, 8]. This notion is supported by some studies that have shown TANs can inhibit metastatic seeding in the lung [9] and that NETosis correlates with better prognosis in ovarian cancer [10]. However, there is much more evidence of the pro-tumoral effects of NETosis. Tumor-promoting NETosis encourages the remodeling of the ECM to promote migration and extravasation, trapping circulating tumor cells in metastasis sites, increasing thrombotic events, increasing secretion of inflammatory factors, protecting tumor cells from other immune cells, and hindering the efficacy of immunotherapies (Figure 1) [6, 11, 12].

**Figure 1:**
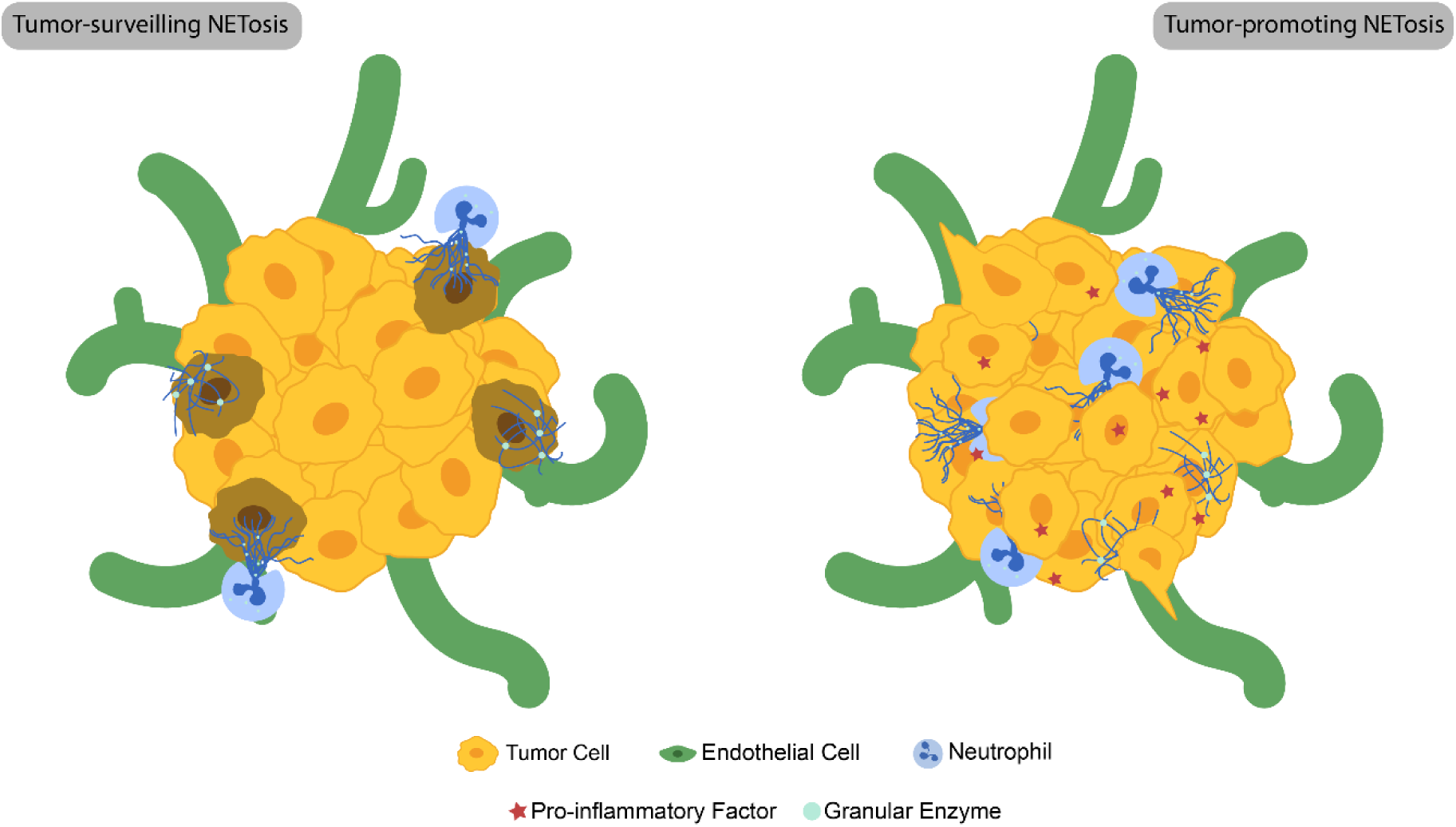
Schematic describing NETosis in cancer. NETs from tumor-surveilling neutrophils in early disease may inhibit tumor progression by trapping cells in the primary tumor with cytotoxic granzymes. NETs from tumor-promoting neutrophils encourage tumor progression by increasing the production of pro-inflammatory factors and encouraging tumor growth, migration and extravasation into the vasculature.

Clinically, both infiltration of TANs in the tumor microenvironment (TME) and high levels of circulating NETs are associated with poor prognosis in most cancers [13, 14]. Recent studies in murine models and 2D culture of human cells have shown that NETs exhibit a chemoattractant effect on triple negative breast carcinoma (TNBC) cells. Yang et al. demonstrated high levels of NETs in the metastatic tissues of patients with advanced TNBC, and that the presence of NETs in distal sites could predict metastasis in a mouse model. Furthermore, they showed that MDA-MB-231 (TNBC cell line derived from a pleural effusion) cell binding to NETDNA was mediated by the transmembrane receptor CCDC25 [13]. Other studies have demonstrated that TNBC cells induce NETosis by secretion of factors like interleukin 8 (IL-8), and that NETosis promotes proliferation, metastasis, and other tumorigenic behavior [12, 15–17].

These murine models have greatly advanced our understanding of the relationship between NETosis and TNBC progression, but there is an opportunity to further probe this relationship in 3D *in vitro* human cell models. There are some emerging studies incorporating neutrophils into TME models in naturally derived hydrogels [3, 18], but the incorporation of NETs alone has been relatively unexplored in 3D *in vitro* models.

The aforementioned work from Yang et al. which established a mechanism by which tumor cells bind directly to NETs implies that NETs can be studied as a matrix biomolecule. Because NETs are large, web-like structures, they can be incorporated into hydrogels via chain entanglement, at which point we hypothesized that cells may interact directly with the NETDNA. In the work presented herein, we developed a reductionist model of NETs in the TNBC microenvironment in a synthetic poly(ethylene glycol) diacrylate (PEGDA)-based hydrogel system we have previously used in TME models [19–22]. To create an enzymatically degradable matrix for to allow for cell migration through the matrix, we inserted GGGPQGIWGQGK (abbreviated as PQ), a matrix metalloproteinase (MMP)-2 and MMP-9 sensitive peptide [23] in the PEGDA backbone, generating PEG-PQ-PEG. We incorporated cell adhesion sites in the hydrogel by conjugating PEG to RGDS, an α_5_β_1_ and α_v_β_3_ integrin binding motif derived from fibronectin [24], and incorporated these dangling PEG-RGDS chains in the hydrogel.

We examined how the incorporation of NETs in PEG-PQ-PEG/PEG-RGDS hydrogels influenced the survival, morphology, proliferation, and MMP secretion of MDA-MB-231 cells. Incorporation of NETs in hydrogels with no cell adhesive sites increased cell survival after 14 days in culture. Addition of NETs to hydrogels increased cell proliferation as measured by the percentage of cells positive for Ki-67 in a dose-independent manner. NETs influenced cell cluster morphology; clusters in gels with high concentrations of NETs showed a more elongated, “migratory” phenotype. NETs also upregulated the secretion of the pro-NETosis factor IL-8. Furthermore, we demonstrated that the conditioned media from MDA-MB-231 cells in gels with NETs induced NETosis in an IL-8-dependent manner.

## Results

### Entrapped NETs Improve Viable Cell Percentage after Long-term Culture in RGDS-free Gels in a Dose-dependent Manner

Adherent cells are known to require integrin activation for long-term survival [25]. While tumor cells of epithelial origin are much more resistant to anchorage-dependent cell death than healthy cells, matrix adherence has been shown to improve survival, increase chemoresistance, and inhibit death-receptor mediated apoptosis [26]. Because tumor cells have been shown to adhere to NET-coated plates [13, 17, 27], we hypothesized that the entrapped NETs would impact adhesion-mediated survival. MDA-MB-231 cells were encapsulated in RGDS-free hydrogels containing varied concentrations of NETs, positive control hydrogels containing 3.5 mM PEG-RGDS, and negative control NET-free, RGDS-free hydrogels. LIVE/DEAD was conducted 14 days after encapsulation, gels were imaged (Figure 2) and the percentage of viable cells was calculated.

**Figure 2:**
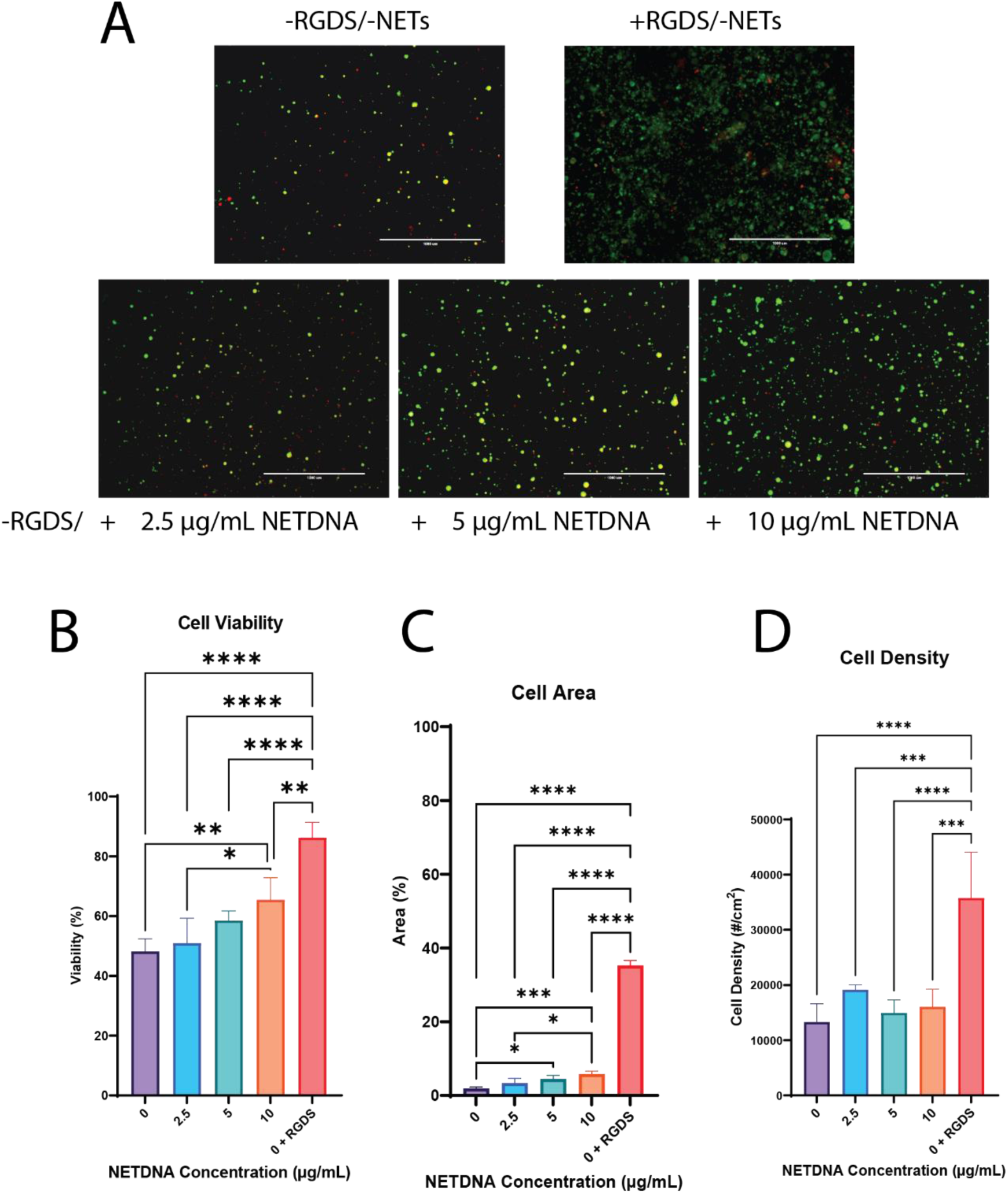
The viability of cells in the gels containing NETs was assessed via LIVE/DEAD assay. A) Representative images of LIVE/DEAD staining in gels. Green is calcein AM (live cells) and red is EthD-1 (dead cells). Scale bars are 1000 µm. B) the percentage of viable cells in the gels. C) The percentage of area occupied by cells. D) The cell density reported as number of cells/cm^2^. The percentage of viable cells and the cell density increased in response to increasing NETDNA concentration, but did not match the levels of the positive control. The cell density increased with the incorporation of PEG-RGDS (mean ± SD, n = 4, * = p < 0.05, ** = p < 0.005, *** = p < 0.0005, **** = p < 0.0001 ANOVA and Tukey’s HSD).

The inclusion of NETs in RGDS-free gels improved cell viability in a dose dependent manner (Figure 2). We also saw a dose-dependent increase in the percentage of area occupied by the cells, implying that the entangled NETs impacted cell spreading (Figure 2). While the cell viability and density increased in response to increased NET concentration, neither approached the cell response seen in the positive control PEG-RGDS gels. Finally, we observed that the inclusion of RGDS significantly increased cell density (Figure 2). The lack of a dose-responsive behavior correlating with the cell area percentage indicates in part that the increase in cell area may largely be due to cell spreading, but also highlights that this method of counting cells may not totally resolve cells that are in close contact.

### Entrapped NETs Increase the Percentage of Proliferating Cells

Because NETosis has been shown to have a positive impact on the proliferation of cancer cells in 2D *in vitro* models as well as in *in vivo* murine models [13, 15, 16], we hypothesized that incorporating NETs into our hydrogels would increase the percentage of proliferating gels. We cultured MDA-MB-231 cells in 3.5 mM PEG-RGDS gels containing 0, 2.5, 5, or 10 µg/mL NETDNA for 3 days prior to fixation and immunostaining for Ki-67. Representative images are shown in Figure 3. We then calculated the percentage of Ki-67 positive cells (Figure 3). Inclusion of NETs in our gels caused a statistically significant increase in the percentage of proliferating cells; however, the concentration of NETs showed no impact. These data suggest that the presence of NETs has a binary, dose-independent (at least within the range studied) impact on tumor cell proliferation.

**Figure 3:**
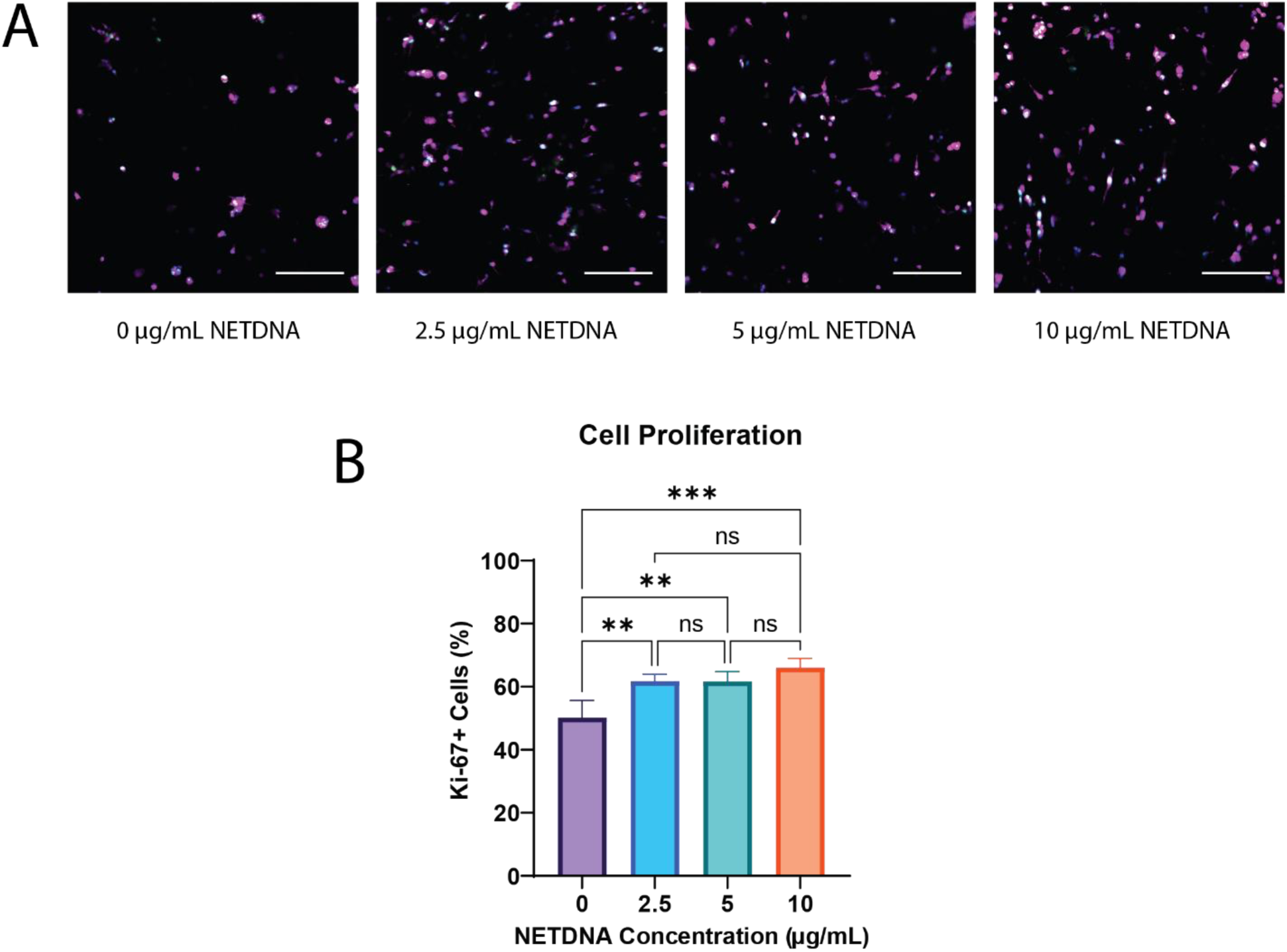
The proliferative activity of cells in gels containing NETs was assessed by immunocytochemistry against Ki-67. A.) Representative images of proliferating cells in gels containing NETs. Pink is phalloidin, blue is DAPI, and green is Ki-67. Scale bars are 200 µm. B). Quantitative analysis of cell proliferation in gels containing NETs. Inclusion of NETs in the gels increased the percentage of Ki-67 positive cells, but NETDNA concentration had no impact (mean ± SD, n = 4, * = p < 0.05, ** = p < 0.005, *** = p < 0.0005, **** = p < 0.0001 ANOVA and Tukey’s HSD).

### Entrapped NETs Influence Cluster Morphology

The adhesion ligand density and composition of a matrix is known to influence cell morphology and cluster formation [28–30]. As such, we hypothesized that the presence of NETs in the synthetic matrix would impact the formation of spheroids in PEG-based hydrogels. We cultured MDA-MB-231 cells in 3.5 mM PEG-RGDS gels containing 0, 2.5, 5, or 10 µg/mL NETDNA for 7 days prior to fixation and staining with DAPI and phalloidin. Representative images are shown in Figure 4. Entrapped NETs impacted both cluster size and circularity in a dose-dependent manner, with higher NETDNA concentrations resulting in distributions trending toward smaller, less circular clusters (Figure 4). Each individual distribution is shown in Supplementary Figures S3 and S4.

**Figure 4:**
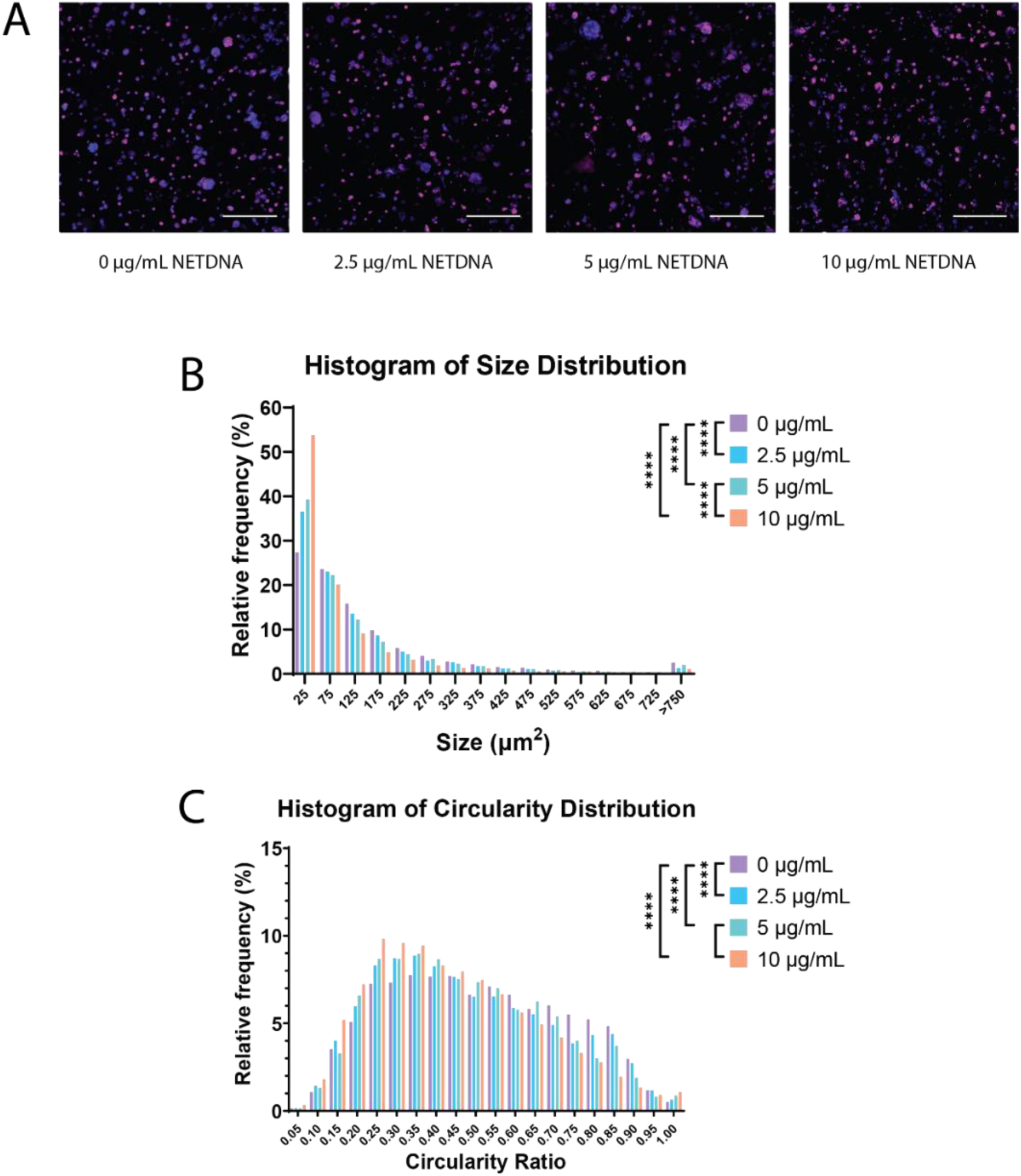
Cluster morphology was visualized by immunocytochemistry. A) Representative images of cell clusters in gels containing NETs. Pink is phalloidin, blue is DAPI. Scale bars are 200 µm. B) Histogram of cell cluster size distribution in gels containing NETs. The relative frequency distributions shifted left (smaller) as NETDNA concentration increased. C) Histogram of cell cluster circularity distribution in gels containing NETs. (n = 4, **** = p < 0.0001 ANOVA and Kruskal-Wallis test with Dunn’s correction).

### Entrapped NETs Influence Secretion of Pro-NETosis Factors

To test whether NETs were capable of inducing production of the pro-NETosis factors IL-8 and IL-18 in a reductionist 3D environment, we cultured MDA-MB-231 cells in 3.5 mM PEG-RGDS gels containing 0, 2.5, 5, or 10 µg/mL NETDNA. We refreshed the media on day 1 and then collected media every 2 days from day 3 to day 9. To examine the change in total secretion levels, we conducted ELISAs for IL-8 and 18, finding minimal IL-18 secretion, and finding by ANOVA that the total IL-8 secretion was dependent on NETDNA concentration in gels (p = 0.0130, Figure 5). The media control had a mean concentration of 923 pg/mL IL-8; all conditioned media samples contained a statistically significant higher concentration than the media control (p < 0.0001). The profiles of IL-8 secretion over time is shown in Supplemental Figure S5.

**Figure 5:**
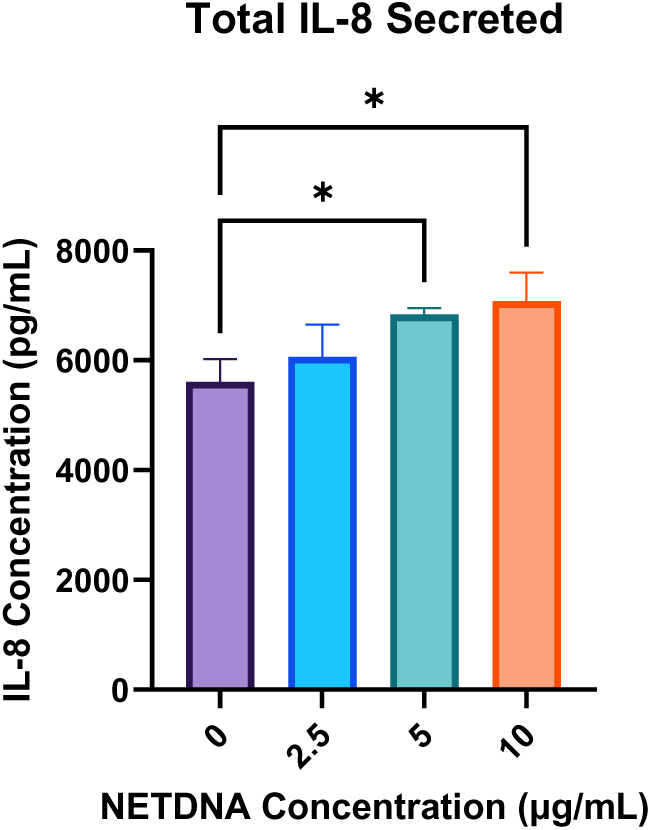
Quantitative analysis of IL-8 secretion. The total amount of IL-8 secreted by cells increased as the NETDNA concentration of gels they were encapsulated in increased (mean ± SD n = 3, * = p < 0.05, ANOVA and Tukey’s HSD).

### Conditioned Media from Cells Cultured in Gels with Entrapped NETs Induces NETosis

Finally, we wanted to determine whether conditioned media from the tumor cell-laden gels was able to induce NETosis in live neutrophils. We cultured neutrophils for 3 hr with phorbol 12-myristate 13 acetate (PMA, stimulates NETosis via the protein kinase C pathway [31]) as a positive control, regular media as a negative control, and conditioned media from each gel type. To half the wells, we added an IL-8 neutralizing antibody. We then fixed, dyed with DAPI and SYTOX GREEN, and stained for citrullinated histone H3 prior to imaging (Figure 6).

**Figure 6:**
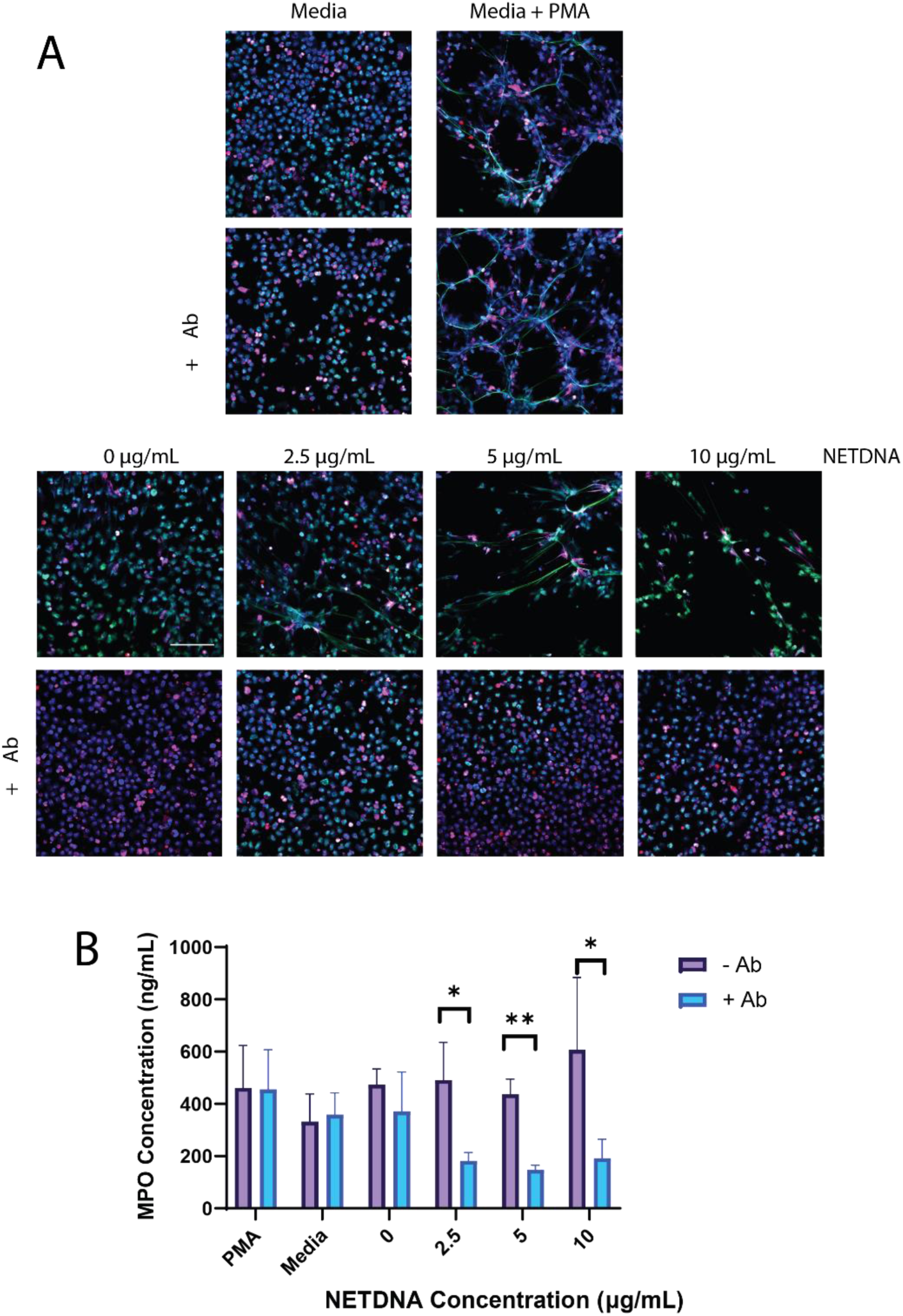
NETosis induced by conditioned media was evaluated via immunocytochemistry and ELISA. A) Images of neutrophils treated with conditioned media from gels containing NETs. Blue is DAPI, green is SYTOX GREEN, pink is citrullinated histone H3. Scale bar is 200 µm. B) Quantitative analysis of MPO concentration in conditioned media. By two-way ANOVA, the IL-8 neutralizing antibody significantly influenced the MPO concentration (p = 0.0003). There was a significant interactive effect between the media type and the addition of the neutralizing antibody (p = 0.0339) Data are presented as mean ± SD (n = 3, abrogation determined by one-tailed t-test).

We quantified the MPO contents of the collected media by ELISA. By two-way ANOVA, we found that the addition of the neutralizing antibody and the interaction between media condition and the addition of the neutralizing antibody significantly influenced the MPO concentration (p = 0.0003 and p = 0.0339 respectively). Specifically, the addition of the IL-8 neutralizing antibody significantly diminished the concentration of MPO in conditioned media from gels containing 2.5 µg/ml (p = 0.0112), 5 µg/mL (p = 0.0006), and 10 µg/mL (p = 0.0330). These data are shown in Figure 6. No MPO was detected in control media samples that had not been cultured on neutrophils.

## Discussion and Conclusions

Recent studies in murine models have made great strides in expanding the understanding of the relationship between the presence of NETs in the TME and tumor progression. NETs have been shown to increase the progression of MDA-MB-231 cells in xenograft tumors, increasing infiltration of distal sites and improving formation of metastases in those distal sites [13]. In 2D culture on tissue culture polystyrene (TCPS), MDA-MB-231 cells have been shown to increase proliferation and pro-NETosis factors [13, 16]. As discussed in Chapter 1, 3D reductionist models are helpful to untangle the influence of individual TME factors in a highly controlled environment that still retains the dimension and physiological stiffness of native tissue. Here, we investigated the influence of entangled NETs in synthetic PEG-PQ-PEG gels.

In order for primary tumors to metastasize, circulating tumor cells must be able to survive without adherence to a matrix, and then in a distal site which may have reduced densities of or different adhesion ligands than in the primary tumor site [32–36]. Since MDA-MB-231 cells have been shown to adhere directly to NETDNA, we hypothesized that NETs could act as an adhesion ligand to promote long-term cell viability in PEG gels with no PEG-RGDS. We demonstrated that the percentage of viable cells as well as the cell density after two weeks increased with higher NETDNA concentrations. The presence of NETs in the liver in a mouse model has been shown to increase TNBC metastasis, and circulating NETs are highly associated with TNBC metastasis in clinic [13, 14]. Our data indicate that a component of these correlations could be due to increased cell survival due to NETDNA binding. The increased cell density in gels with higher NETDNA concentration also suggested that NETs can increase proliferation in low-adhesive environments.

Progression of both primary tumors and metastases is dependent on the ability of cells to grow and proliferate. In Yang et al., MDA-MB-231 cells cultured in 2D on TCPS proliferated more in response to treatment with NETs [13]. They demonstrated that this behavior was adherence-dependent, as cells with CCDC25 knocked out did not show the same increase in proliferation. Thus, we hypothesized that cells cultured in PEG-PQ-PEG/PEG-RGDS gels with NETs would show an increase in Ki-67 expression, showing cells in the active phase of the cell cycle. We found that the incorporation of NETs did increase the percentage of proliferating cells, but this did not depend on NETDNA concentration. The impact of NETs on proliferation in this 3D environment correlates with the NETs frequently observed in metastasis sites; perhaps the NETs promote metastasis formation not only by acting as a chemoattractant, but also by increasing proliferation once tumor cells colonize the site.

Because adhesion ligands are known to influence cluster and spheroid morphology [21], we wanted to investigate the influence of entrapped NETs on cluster size and circularity. We observed that as NETDNA concentration increased, clusters became smaller and less spherical. The decrease in size and circularity reflects a more migratory, pro-metastatic phenotype, which is consistent with *in vivo* studies that found NETs increased the metastatic behavior of MDA-MB-231 cells [13, 16].

Many cytokines induce NETosis, but IL-8 is one of particular interest in tumor-associated NETosis. It has been demonstrated to be secreted by many tumor cell types, including triple negative breast cancer, and tumor-secreted IL-8 has been shown to induce NETosis [12, 17, 37]. A feedback loop in which the presence of NETs induced IL-8 secretion by colorectal cancer cells, which then induced NETosis, was established in a murine model [17]. Additionally, IL-18 has been shown to be upregulated in in tumor cells, specifically the MDA-MB-231 cell line [38, 39], and the presence of NETs has been shown to upregulate IL-18 production in wound healing as well as several other diseases [40, 41]. These factors both represent components of potential positive feedback loops in tumor-associated NETosis: the presence of NETs increases the production of IL-8 and 18 by tumor cells; secreted IL-8 and 18 induce NETosis, resulting in an increased number of NETs in circulation and in the TME. We did not observe significant IL-18 expression in MDA-MB-231 cells cultured in PEG-PQ-PEG/PEG-RGDS gels containing NETs. However, we observed that IL-8 secretion was dependent on NETDNA concentration. This result indicates that the presence of NETs in the microenvironment model influences the secretion profile of a known pro-NETosis factor.

After demonstrating that entrapped NETs influenced IL-8 secretion, when investigated the ability of conditioned media to influence NETosis. By immunocytochemistry, we observed that conditioned media from cells cultured in gels with entrapped NETs was able to stimulate NETosis, and that this effect appeared to be abrogated by the addition of an IL-8 neutralizing antibody. We quantified the amount of MPO in the media collected from the stimulated neutrophils, finding that the MPO concentration was reduced by the addition of the IL-8 neutralizing antibody in the 2.5 µg/ml, 5 µg/mL, and 10 µg/mL NETDNA conditions. We did find MPO even in conditions were significant NETosis was not observed by immunocytochemistry, indicating that the media in those conditions was directing neutrophils toward an active, MPO-secreting state [42, 43]. These results, together with the IL-8 secretion results, indicate a feedback loop in our model in which NETs influence cells to produce more IL-8, which in turn stimulates NETosis. These results are similar to the feedback loop established in an *in vivo* colorectal cancer model [17]. These data help link together previous 2D *in vitro* studies that indicated that NETs induce IL-8 secretion in MDA-MB-231 cells [16] and that this conditioned media can induce NETosis in an IL-8 dependent manner [44], and *in vivo* studies that indicated that MDA-MB-231 cells can stimulate NETosis [37].

In this work, we demonstrated that entangled NETs in PEG-PQ-PEG gels influence MDA-MB-231 progression in several ways. NETs increase cell viability in the absence of other adhesion ligands, increase cell proliferation, increased matrix degradation, and cause smaller, more elongated spheroid formation. All of these behaviors correlate with more advanced, more metastatic tumors. Furthermore, NETs influenced the secretion profile of IL-8, a pro-NETosis factor. Conditioned media from cells in gels containing NETs influenced NETosis in an IL-8-dependent manner. We developed a 3D, reductionist model suitable for isolating the influence of NETs on tumor progression.

## Methods

### Cell Maintenance

MDA-MB-231 (ATCC, Manassas, VA) cells and primary human neutrophils (Hemacare, Los Angeles, CA) were cultured in MEM-α (Thermofisher Scientific, Waltham, MA) with fetal bovine serum (10%, Sigma-Aldrich, St. Louis, MO) and penicillin-streptomycin (1%, VWR, Radnor, PA). Media were refreshed every 2-3 days. Cells were detached for passaging by 5 min incubation with trypsin-EDTA (0.25%, VWR). MDA-MB-231 cells were used in studies between passages 45 and 50. Primary human neutrophils were used directly out of thaw and cultured on poly-l-lysine (15 µg/mL, Thermofisher) coated tissue culture dishes. All cells were maintained at 37 °C and 5% CO_2_.

### NET Extraction

NETosis was induced in primary human neutrophils using a protocol adapted from Najmeh et al. [27] Briefly, NETs were incubated in media containing 5 mM PMA (Sigma-Aldrich) for 4 hr. Media was gently aspirated, and the neutrophils and NETs were collected by repeated rinsing of the flask in cold DNAse and RNAse free ultrapure water (VWR) before centrifugation at a speed of 450 xg for 10 min to remove neutrophils. The supernatant was collected, and NETs were concentrated by sequential centrifugation steps at a speed 17,000 xg for 10 min at 4 °C. The final concentration of NETs in terms of both NETDNA and entrapped proteins was measured on a NanoDrop2000 (Thermofisher) using the nucleic acid 260/280 function and protein 280 and 205 functions. The nucleic acid function uses the absorbance at 260 nm to measure nucleic acid content, and the absorbance at 280 nm to measure contaminating protein content [45]. The NET stocks were stored at 4 °C. To be consistent with literature, we entrapped NETs based on the DNA concentration.

### Boyden Chamber Assay

In order to confirm that our MDA-MB-231 cells respond to our extracted NETs in a manner similar to what has been seen in literature, we conducted a dose-response Transwell migration experiment similar to the supplemental study from Yang et al. [13] MDA-MB-231 cells were treated with 25 µg/mL mitomycin C (Sigma-Aldrich) for 2 hr to inhibit proliferation, then seeded onto fibronectin-coated 8 µm Transwell inserts (Corning) at a density of 10,000 cells/insert. Cells were allowed to migrate toward media with 0, 1, 2, or 5 µg/mL NETs for 4 hr prior to fixation with 4% paraformaldehyde (Electron Microscopy Services, Hatfield, PA) in phosphate-buffered saline (PBS, Sigma-Aldrich) for 15 min at room temperature and staining with 0.005% Crystal Violet in PBS (Sigma-Aldrich) for 10 min at room temperature. The inside of the inserts was wiped with a cotton swab to remove any unmigrated cells before taking phase contrast images on a Zeiss Axiovert 135 epifluorescence microscope (10X Plan-NEOFLUAR objective, NA: 0.3, Zeiss, Oberkochen, Germany). The number of migrated cells per field of view was counted manually using the multipoint tool in ImageJ (NIH). These results are shown in the Supplementary Figure S1.

### Conjugation of RGDS and PQ to PEG and PEG-Peptide Characterization

PQ and RGDS were conjugated to PEG as described previously [19]. To generate cell-adhesive PEG-RGDS, 3.2 kDa acrylate-PEG-succinimidyl valerate (acryl-PEG-SVA, Laysan Bio, Arab, AL), RGDS peptide (Genscript Biotech, Piscataway, NJ), and diisopropylethylamine (DIPEA, Sigma-Aldrich) were dissolved in anhydrous dimethyl sulfoxide (DMSO, Sigma-Aldrich) and reacted overnight at room temperature under inert gas. The ratio of DIPEA:PEG-SVA was 2:1 and the ratio of RGDS:PEG-SVA was 1.2:1.

To generate enzymatically degradable PEG-PQ-PEG, similar methods were used with GGGPQGIWGQGK (Genscript), abbreviated as PQ, in place of RGDS. Because the PEG was conjugated to both the N-terminal amine group and the amine in the side group of the C-terminal lysine, the reaction ratio of PQ:PEG-SVA was adjusted to 1:2.

The products were then dialyzed using 3.5 kDa molecular weight cutoff (MWCO) dialysis membrane (Thermofisher) for PEG-RGDS and 6-8 kDa MWCO dialysis membrane for PEG-PQ-PEG (Thermofisher), then lyophilized. They were then dissolved in dimethylformamide with 0.1% ammonium acetate and evaluated by gel permeation chromatography (GPC) with an evaporative light scattering detector. Plots of PEG-peptide column retention time were compared against that of unconjugated Acryl-PEG-SVA. The larger a sample is, the shorter the retention time. Peaks corresponding to PEG-PQ-PEG and PEG-RGDS that occurred earlier than the peak corresponding to PEG-SVA indicated successful conjugation. Any unreacted PEG-SVA in a PEG-peptide sample generates a secondary peak that cooccurs with the PEG-SVA sample peak. All products had conjugation efficiencies above 90%.

### Encapsulation of Cells in Hydrogels containing NETs

First, glass slides were rendered hydrophobic via treatment with Sigmacote (Sigma-Aldrich). The slides were rinsed with ultrapure water followed by acetone, submerged in Sigmacote, then air-dried. This process was repeated 5 times. Next, methacrylate groups were added to the surface of coverglass. The coverglass was etched in Piranha (30% hydrogen peroxide (EMD Millipore, Burlington, MA, USA) + 70% sulfuric acid (VWR)) for 1 hr, then reacted for 4 days in 95% ethanol (Decon Labs, King of Prussia, PA, USA) with 2 v*/*v% 3-(trimethoxysilyl) propyl methacrylate) (TMSPA, Sigma-Aldrich). These modifications allowed for easy manipulation of the crosslinked hydrogels: the hydrophobic glass slides do not interact with the hydrophilic hydrogels, while the methacylate-modified coverglass attaches to the hydrogel during crosslinking, allowing the coverglass and attached hydrogel to be removed from the glass slide with forceps.

To create hydrogels with cells encapsulated in 3D, 3 w/v% PEG-PQ-PEG and 3.5 mM PEG-RGDS were dissolved in HEPES-buffered saline (HBS, pH 8.3) with 1.5% (v/v) triethanolamine (TEOA, Sigma-Aldrich), 0.01 mM eosin Y, and 0.35% (v/v) N-vinylpyrrolidone (NVP, Sigma-Aldrich) for photoinitiation of crosslinking. All materials were sterilized by filtration through a 0.2 μm syringe filter. NET stock was added to the polymer precursor solution to the desired final concentration of NETDNA. The precursor solution was then vortexed on the highest speed for 45 sec to mix. Cells were passaged, centrifuged, and the pellet was resuspended in the polymer precursor solution. A 5µL droplet of the cells suspended in hydrogel precursor solution was placed on a Sigmacote-modified glass slide between two 380 µm PDMS spacers. A 5µL droplet of the precursor solution was placed on a Sigmacote-treated glass slide between two 380 µm polydimethyl siloxane (PDMS) spacers. A methacrylate-modified coverglass was placed on the spacers, compressing the droplet into a cylinder which was 380 µm tall and roughly 3.8 mm in diameter. The gel was crosslinked upon exposure to white light at 280 mW/cm^2^ for 40 sec. The methacrylate-modified coverglass with the attached gel was removed and placed in a 24-well plate. Gels were covered with the cell culture media and transferred to a tissue-culture incubator at 37°C and 5% CO_2_ for cell maintenance. A schematic of this process is shown in the Supplementary Figure S2.

For the proliferation and morphology studies, cells were encapsulated at a density of 3 x 10^6^ cells/mL. For the LIVE/DEAD assay, cells were encapsulated at a density of 2 x 10^6^ cells/mL, because a higher density resulted in degradation of the high-NET content gels before the 14 day time point. For IL-8 & 18 secretion studies, cells were encapsulated at a density of 5 x 10^6^ cells/mL to help increase the concentration of factors for analysis.

### Immunocytochemistry

Staining was conducted in PBS. All samples were fixed with 4% paraformaldehyde (Electron Microscopy Services) for 45 min. For the proliferation study, cells were fixed after 3 days in culture. For the morphology study, cells were fixed after 7 days inn culture. Cell membranes were permeabilized via incubation with 0.25% Triton-X 100 (Sigma-Aldrich) for 45 min, rinsed 4 times in PBS, and blocked overnight at 4 °C in 5% donkey serum (Sigma-Aldrich). For the proliferation study, the samples were incubated with a rabbit-anti-Ki-67 primary antibody (1:1000, Abcam) in 0.5% donkey serum for 48 hr at 4 °C. Samples were then rinsed with 0.01% Tween20 (Sigma-Aldrich) three times, followed by one rinse in PBS over a period of 8 hr. The samples were then incubated with an Alexafluor 488 donkey-anti-rabbit secondary antibody (1:200, Thermofisher) in 0.5% donkey serum for 48 hr at 4 °C. Samples were then rinsed in PBS for 2 hr. All samples were then incubated with 4′,6-Diamidino-2-phenylindole (DAPI, 2 µM, Thermofisher) as a nuclear counterstain and Alexafluor 555 phalloidin (1:60, Thermofisher) as an F-actin counterstain for 1 hr at 4 °C. Samples for the morphology study were stained only with DAPI and phalloidin to visualize cell shape.

### LIVE/DEAD Assay

Cells were encapsulated in gels containing no NETs or PEG-RGDS (negative control), gels with no PEG-RGDS and 2.5, 5, or 10 µg/mL of NETDNA, or gels with no NETs and 3.5 mM PEG-RGDS (positive control) and cultured for 14 days. Cells were then treated with the LIVE/DEAD Viability/Cytotoxicity kit (Thermofisher) according to the manufacturer’s protocol. The gels were immediately imaged on an EVOS FL Auto microscope (4X LPlan PH2 objective, NA: 0.13, Life Technologies, Carlsbad, CA).

The percentage of viable cells remaining in the gels after 14 days was determined using a macro created in ImageJ (NIH, Bethesda, MD). Prior to running the macro, the red and green channels were separated, and images were converted to 8-bit. In brief, the backgrounds of the images were subtracted, and the local contrast was enhanced using the CLAHE plugin. The image was then converted from grayscale to binary black and white using the threshold command. Finally, the analyze particles command was executed using a minimum size threshold of 10 µm^2^ to exclude any possible artifacts below the expected size of a nucleus, resulting in an output of the approximate number of cells in the image, as well as an area percentage of the image covered by cells. The percentage of viable cells remaining was calculated as 100 times the number of cells in the green channel divided by the sum of the number of cells in the green channel and the number of cells in the red channel. The cell density was calculated as the total number of cells divided by the area of the image.

### Quantitative Analysis of Cell Proliferation

Images were taken on an 880 Inverted AiryScan microscope (10X Plan-NEOFLUAR objective, NA: 0.3, Zeiss) with 2.4 µm slice thickness. 30 µm Z-projections were used for analysis, with 4 fields of view per gel. Ki-67 positive and negative nuclei were counted manually using the cell counter function in ImageJ (NIH). The percentage of proliferating cells was calculated as 100 times the number of Ki-67 positive nuclei divided by the total number of nuclei.

### Quantitative Analysis of Cell Cluster Morphology

Images were taken on an 880 Inverted AiryScan microscope (10X Plan-NEOFLUAR objective, NA: 0.3, Zeiss) with 2.4 µm slice thickness. 100 µm Z-projections were used for analysis, with 4 fields of view per gel. Cluster morphology was analyzed using a macro developed in ImageJ (NIH). In brief, images were converted to RGB color to collapse the DAPI and phalloidin channels into one image, then converted to 8-bit. Grayscale images were converted to binary black and white using the threshold command. The analyze particles function was run with a minimum size of 10 µm^2^ to exclude any staining artifacts below the size of a nucleus, resulting in an output of the size and circularity of each cell cluster.

### Secreted Factors Study

For the IL-8 and IL-18 secretion study, cells were encapsulated as described in section 5.2.4, but in larger 20 µL gels with 0, 2.5, 5, or 10 µg/mL of NETDNA. The larger gels serve to increase the concentration of secreted factors in media samples, allowing for easier detection. On day 1, the media was aspirated and replaced. For the multiple timepoint study, cells were cultured in 800 µL media per well. Media was collected and replaced every 2 days up until day 9. All media samples were stored at −80 °C for later analysis. For the conditioned media study, media was refreshed on day 1 and then collected day 7.

The following enzyme-linked immunoassay (ELISA) kits were used to analyze secreted factors: IL-18 (BMS267-2, Thermofisher) and IL-8 (D8000C, R&D Systems). Samples were tested at 1:100 dilutions. ELISAs were run according to the manufacturers’ protocols.

### Conditioned Media NETosis Study

Neutrophils were seeded on poly-l-lysine (15 µg/mL, Thermofisher) coated Labtek 8-well chamber slides (Thermofisher) at a density of 360,000 cells/cm^2^. They were treated with media containing 5 mM PMA (positive control), normal media (negative control), or reserved conditioned media from cells encapsulated in gels with varying concentrations of NETs (Section 5.2.9). An IL-8 neutralizing antibody previously used by Yamato et al. [46] (15 µg/mL, AHC0881, Thermofisher) was added to half the wells.

After 3 hr, the media was collected and stored at −80 °C for later analysis, and cells were fixed in 4% paraformaldehyde (Electron Microscopy Services) for 15 min at room temperature. Cells were rinsed with PBS three times, then blocked in 5% donkey serum for 1 hr at 4 °C. Cells were incubated with rabbit-anti-citrullinated histone H3 primary antibody (1:1000, Abcam) in 0.5% donkey serum for 4 hr at 4 °C. Samples were again blocked in 5% donkey serum for 1 hr at 4 °C, then incubated with Alexafluor 555 donkey-anti-rabbit secondary antibody (1:200, Thermofisher) in 0.5% donkey serum overnight at 4 °C. Samples were rinsed three times in in PBS, then stained with DAPI (2 µM, Thermofisher) as a permeant nuclear counterstain and SYTOX GREEN (2 drops/10mL, Thermofisher) as a non-permeant nuclear counterstain, for 30 min at 4 °C. Samples were rinsed and stored in PBS.

Images were taken on an 880 Inverted AiryScan microscope (10X Plan-NEOFLUAR objective, NA: 0.3, Zeiss). The MPO content of the collected media was analyzed by ELISA (1:100 dilution, DMYE00B, R&D Systems) according to the manufacturer’s protocol.

### Statistical Analysis

All statistical analyses were performed in GraphPad Prism 9 (Graphpad Software, Inc, San Diego, CA). The LIVE/DEAD and proliferation results were analyzed by ANOVA. The statistical difference between each pair was determined via ANOVA with Tukey’s Honest Significant Difference (HSD) post-hoc test to allow for comparison across multiple groups. The morphology results were analyzed by the Kruskal-Wallis ANOVA test for comparing distributions, and the statistical difference between each pair was determined with the Dunn’s multiple comparisons post-hoc correction. The IL-8 ELISA data was analyzed by ANOVA to determine the impact of NET concentrations on the total amount of IL-8 secreted. Pairwise comparisons were determined by ANOVA with Tukey’s HSD. The MPO ELISA data was analyzed by two-way ANOVA to determine the impact of conditioned media type and antibody neutralization. To determine whether the addition of the neutralizing antibody significantly diminished MPO concentration for a given condition, data were analyzed by one-tailed Student’s t test. All *p-*values less than 0.05 were considered statistically significant.

## Acknowledgements

This work was funded by the Duke Graduate School and the NIH T32 Training Grant. The authors would like to thank Prof. Joel Collier and Prof. Daniel Reker for providing laboratory space to conduct this work.

## Author Contributions

JLW conceived of the project. RRK wrote the manuscript, designed and executed experiments, and conducted data analysis. SS conducted ELISA experiments and assisted with ELISA data analysis. JLW supervised the project and edited the manuscript.

## Data Availability Statement

The datasets generated during and/or analyzed during the current study are available from the corresponding author on reasonable request.

## Competing Interests Statement

The authors declare no competing interests.

## Supplementary Information

### Boyden Chamber Assay

We conducted a preliminary NET dose-response experiment to confirm that we were seeing the same chemoattractant properties reported in literature. MDA-MB-231 cells pre-treated with mitomycin C to inhibit proliferation were seeded atop Transwell inserts with 8 μm pores. Media with NET concentrations varying from 0 to 5 μg/mL was added to the bottom of the well, and cells were allowed to migrate for 4 hours prior to fixation. Cells on the underside of the membrane were stained with Crystal Violet. We saw a dose-dependent increase in migrated cells per field of view, as has been previously established in literature (Figure 29).

**Figure S1:**
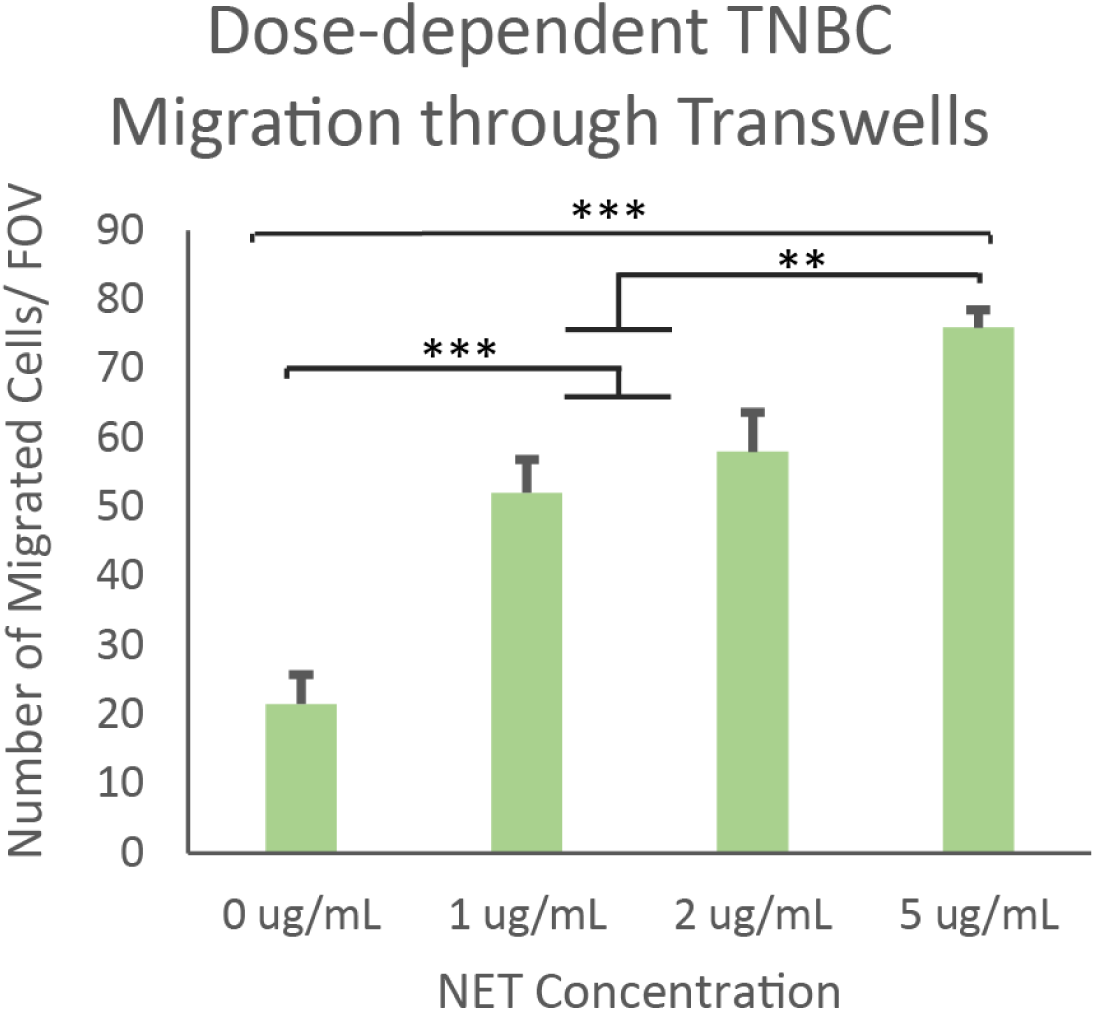
Migration of MDA-MB-231 cells through a Transwell membrane in response to NET concentration. (mean ± SD, n = 3, * = p < 0.05, ** = p < 0.005, *** = p < 0.0005, ANOVA and Tukey’s HSD)

### NET Entanglement in PEG Gels

**Figure S2:**
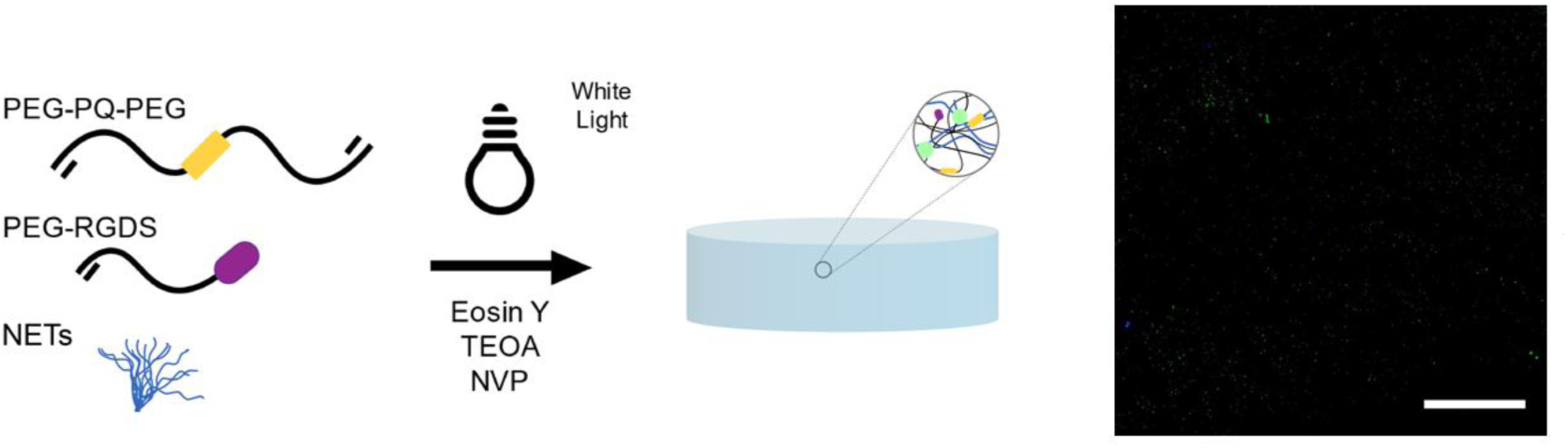
Entanglement of NETs in PEG gels. NETs were incorporated into the polymer precursor solution by vortexing on high speed for 45 sec prior to polymerization under white light. Punctate MPO staining (green) was observed in gels with entrapped NETs. Scale bar = 200 µm.

### Cell Cluster Morphology

**Figure S3:**
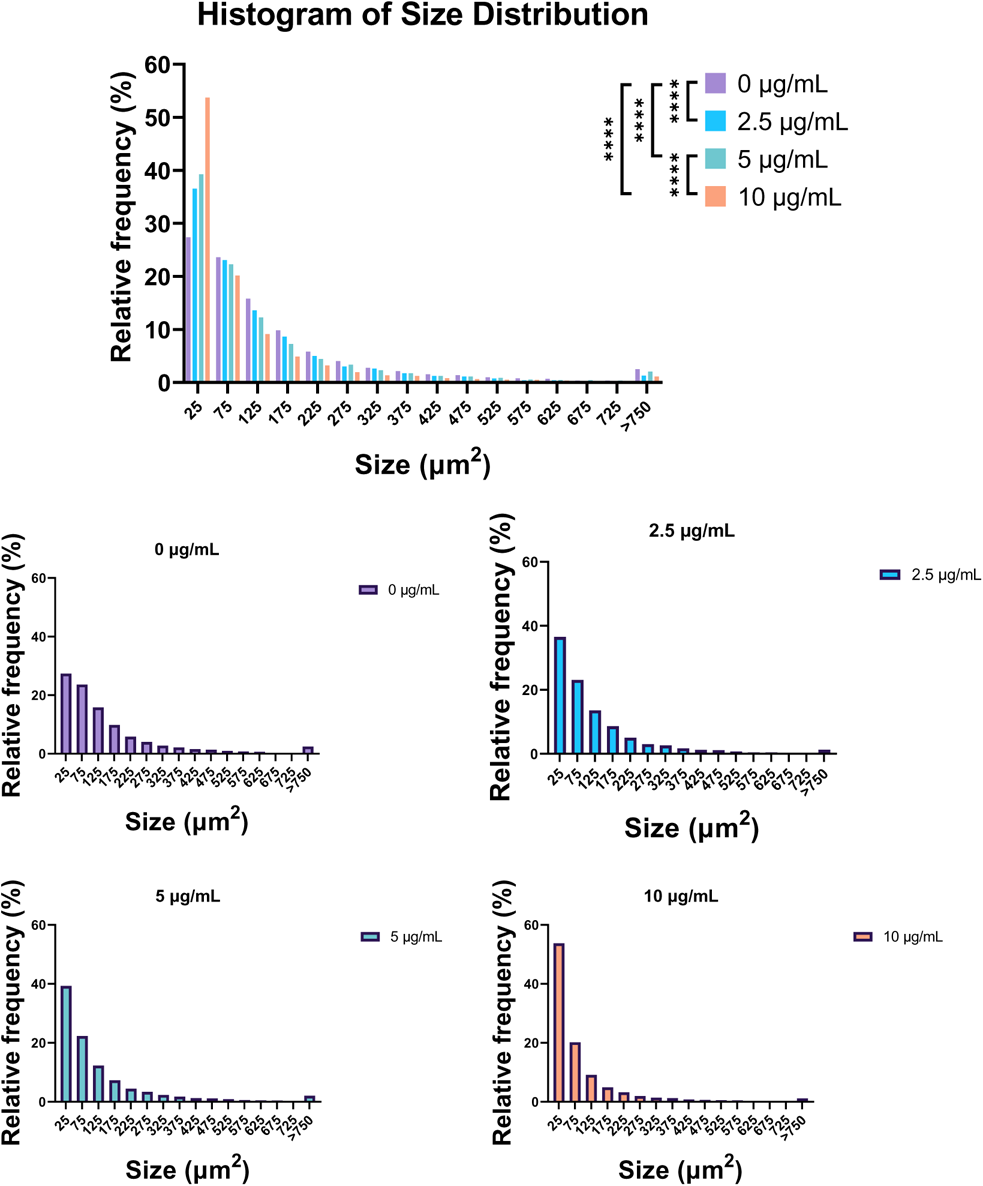
Histograms of cell cluster size distribution in gels containing NETs. Top: distributions plotted on one graph, bottom: distributions separated by NETDNA concentration. The relative frequency distributions shifted left (smaller) as NETDNA concentration increased. (**** = p < 0.0001 ANOVA and Kruskal-Wallis test).

**Figure S4:**
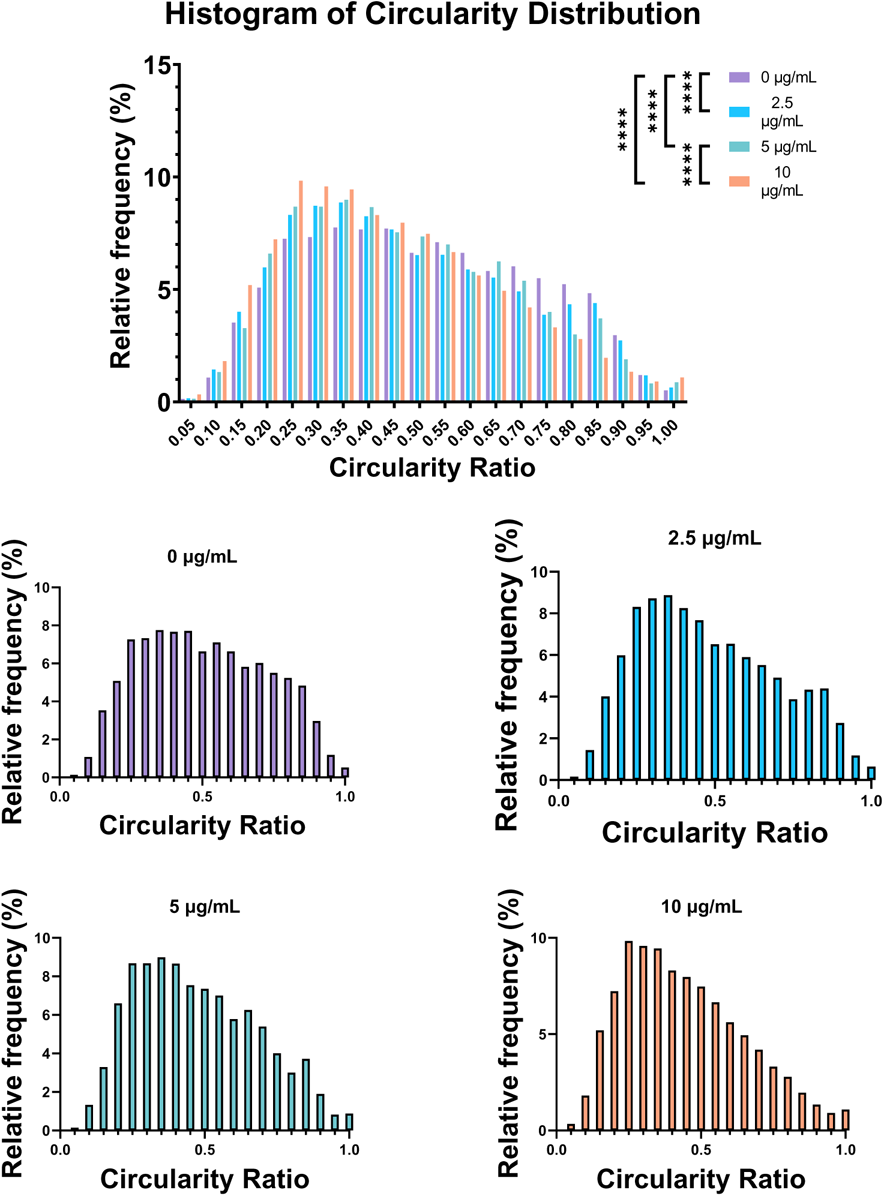
Histograms of cell cluster circularity distribution in gels containing NETs. Top: distributions plotted on one graph, bottom: distributions separated by NETDNA concentration. The relative frequency distributions shifted left (less circular) as NETDNA concentration increased. (**** = p < 0.0001 ANOVA and Kruskal-Wallis test).

### IL-8 Secretion Over Time

**Figure S5:**
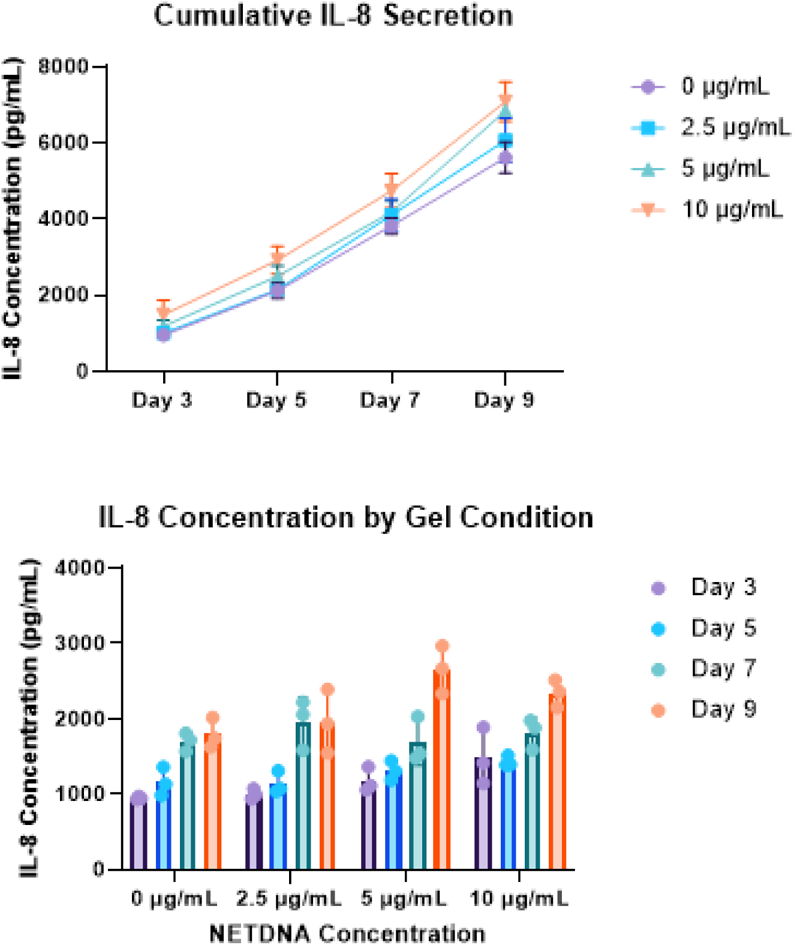
Quantitative analysis of IL-8 secretion over time. Top: cumulative IL-8 secretion by cells over time increased as the NETDNA concentration of gels they were encapsulated in increased. Bottom: bar chart showing IL-8 secretion at each time of collection. By two-way ANOVA, NETDNA concentration (p = 0.013) and day of collection (p < 0.0001) impacted IL-8 secretion. Data are presented as mean ± SD (n = 3).

#### Influence of Entrapped NETs on Matrix Degradation

Matrix remodeling is a hallmark of tumor progression and is required for growth and metastasis. We hypothesized adding NETs to the synthetic matrices would impact the change in matrix stiffness over time. Cell-laden gels containing 0, 2.5, 5, or 10 µg/mL of NETDNA were fabricated. For mechanical testing, gels were cultured for 7 days. Media was aspirated and replaced with PBS, then samples were immediately subjected to compressive testing as described previously (Katz and West, 2022). To visualize macroscopic changes in gel appearance, gels were photographed on day 1, and then every time media was refreshed over a period of 9 days. Images were taken on an Iphone 11 (Apple, Cupertino, CA).

We observed that the gels with higher NET contents appeared “softer,” with less defined edges over time (Figure S6). To quantify this change in stiffness, we cultured MDA-MB-231 cells in the same four gel conditions for 7 days prior to subjecting them to compression testing. In the 10 µg/mL condition, gels were so soft that they deteriorated prior to reaching the linear region of the stress-strain curve, and no data could be collected.

**Figure S6:**
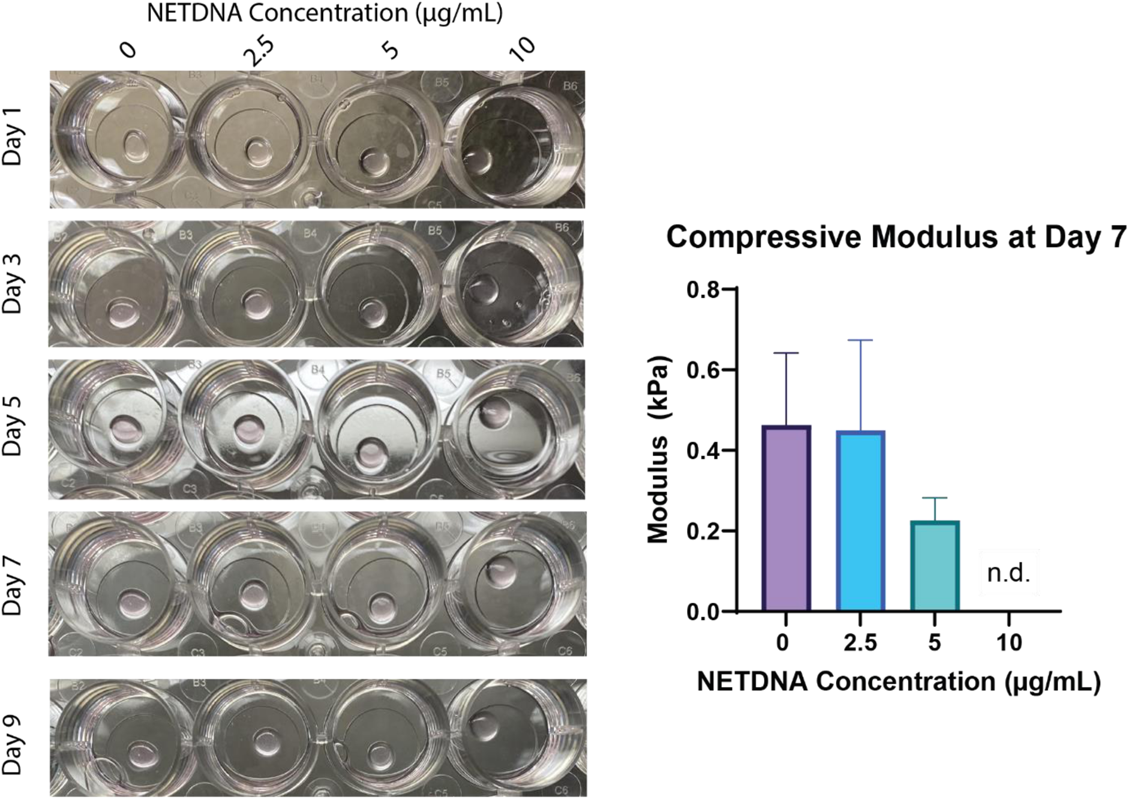
NET concentration impacts gel softening. Left: images of change in gel structure over time. The crisper edge seen in all gels on Day 1 disappears most rapidly in the 10 µg/mL NETDNA condition. Right: compressive moduli of gels with varied concentrations of NETDNA at Day 7. Data are presented as mean ± SD (n = 3).

In addition to compressive testing, we also assessed MMP secretion. Cells were encapsulated in gels with 0, 2.5, 5, or 10 µg/mL of NETDNA. On day 1, the media was aspirated and replaced. For the multiple timepoint study, cells were cultured in 400 µL media per well. Media was collected and replaced every 2 days up until day 9. For the weeklong study, 800 µL media was added per well and collected on day 7. Collected media was tested at a 1:2 dilution using MMP-2 (KHC3081, Thermofisher) and MMP-9 (DMP900, RND Systems) ELISA kits.

We found that MMP-2 secretion was independent of NETDNA concentration. However, trends in MMP-9 secretion indicated a possible dependence on NETDNA concentration. Specifically, it seemed that cells cultured in gels with a higher concentration of NETDNA secreted more MMP-9 earlier on cell culture, resulting in more rapid degradation of the gels (Figure S7).

**Figure S7:**
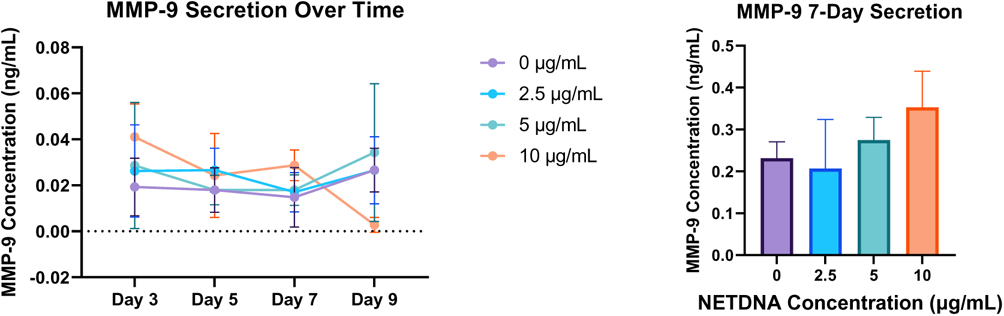
Quantification of MMP-9 secretion by ELISA. Trends indicate the possibility of increased secretion caused by NET entrapment in gels, but further investigation is required. Data are presented as mean ± SD (n = 3).

These data were not statistically significant, and thus further investigation is required to determine the mechanism by which NETs increase matrix degradation.

## Notes

### Competing Interest Statement

The authors have declared no competing interest.

